# Multiparametric MRI and imaging transcriptomics reveal molecular and cellular correlates of neurodegeneration in experimental parkinsonism

**DOI:** 10.1101/2025.11.12.688033

**Authors:** Eugene Kim, Diana Cash, Daniel Martins, Camilla Simmons, Antonio Heras-Garvin, Elena Klippel, Florian Krismer, Laura Mantoan Ritter, Nadia Stefanova

## Abstract

Multiple system atrophy (MSA) is an atypical Parkinsonian disorder marked by oligodendroglial α-synucleinopathy and selective neurodegeneration. Although MRI can capture regional atrophy and microstructural alterations in MSA brain, the molecular substrates underlying these phenotypes remain poorly defined. Imaging transcriptomics provides a computational framework to relate spatial imaging patterns to brain-wide gene expression. While this approach has been applied to human MSA, interpretation is constrained by limited experimental control and lack of disease-matched molecular validation. Here, we apply imaging transcriptomics in a controlled preclinical setting by integrating high-resolution ex vivo multimodal MRI with transcriptomic mapping in the PLP-αSyn mouse model of MSA. Structural and diffusion MRI revealed distinct patterns of regional atrophy and microstructural abnormalities. Atlas-based analyses associated imaging phenotypes to gene programs related to oligodendrocyte biology, energy metabolism, and neuroinflammation, with modality-specific signatures. These associations were supported by independent RNA-sequencing and showed convergence with human MSA findings. Our work benchmarks MRI–transcriptomic relationships in MSA and provides a translational framework for interpreting imaging biomarkers in synucleinopathies.

## INTRODUCTION

Multiple system atrophy (MSA) is a rapidly progressive neurodegenerative disorder clinically characterized by autonomic failure, parkinsonism, and cerebellar ataxia. Its etiology remains unclear, diagnosis is challenging, and no effective disease-modifying therapy is currently available [1]. MSA is classified as an α-synucleinopathy due to the hallmark accumulation of α-synuclein (α-syn) fibrils within oligodendrocytes, forming glial cytoplasmic inclusions (GCIs) [2]. At the systems level, MSA is characterized by regionally selective vulnerability that likely reflects complex interactions across molecular-, cellular-, and network-scale processes. Magnetic resonance imaging (MRI) provides a powerful, non-invasive window into these processes by capturing macroscopic and microstructural brain alterations in vivo (**Table 1**). However, while MRI phenotypes such as atrophy or diffusion abnormalities are sensitive markers of disease progression [3, 4], they lack specificity and the molecular and cellular substrates underlying these imaging changes remain incompletely characterized.

**Table 1:** Regions of interest with significantly altered MRI volumes and diffusivity in human MSA compared to healthy controls.

| Brain region | MRI |  |  |  |
| --- | --- | --- | --- | --- |
|  | volume | FA | MD | Ref. |
| Cerebral white matter | ↑ | ↓ | ↑ | [36-41] |
| Amygdala | - | ↓ | - | [42] |
| Pyramidal tract | ↓ | ↓ | ↑ | [37, 43-49] |
| Capsula interna | - | ↓ | ↑ | [36-38, 43, 45, 50, 51] |
| Putamen/Striatum | ↓ | ↓ | ↑ | [36, 42, 44, 52-62] |
| Globus pallidus | - | ↑ | - | [42, 60] |
| Midbrain | ↓ | ↓ | ↑ | [38, 58, 60, 63, 64] |
| Cerebral peduncle |  | ↓ | ↑ | [38, 51] |
| Substantia nigra | ↓ | ↓ | - | [42, 56, 58, 65, 66] |
| Red nucleus | - | - | ↑ | [42] |
| Subthalamic nucleus | - | - | ↑ | [42] |
| Brainstem | - | - | ↑ | [67] |
| Medulla | ↓ | - | - | [56] |
| Pons | ↓ | ↓ | ↑ | [37, 38, 44, 45, 48, 50, 51, 56, 58-61, 63, 65, 68-72] |
| Cerebellum | gray matter: ↓<br>white matter: ↑/↓<br>total: ↓ | ↓ | ↑ | [38-42, 44-46, 48, 52, 53, 56, 58-61, 67, 70, 71, 73, 74] |
| Middle cerebellar peduncle (MCP) | ↓ | ↓ | ↑ | [36-38, 42, 44-46, 48-50, 56, 58-61, 68, 70, 75-79] |
| Superior cerebellar peduncle (SCP) | - | ↓ | ↑ | [36-38, 60, 78] |

↑ = significantly increased ↓ = significantly decreased - = no data available FA = fractional anisotropy MD = mean diffusivity

To address this limitation, imaging transcriptomics has emerged as a framework for linking spatial patterns of neuroimaging phenotypes to normative brain-wide gene expression maps derived from resources such as the Allen Brain Atlas. Rather than measuring disease-state transcription directly, imaging transcriptomics exploits the correspondence between the spatial organization of normative gene expression and the regional distribution of disease-related imaging alterations to identify molecular and cellular programs associated with regional vulnerability across the brain. This approach has been applied extensively in humans to interpret MRI-derived cortical thinning, volumetric atrophy, and functional alterations in neurodegenerative disorders, including Alzheimer’s disease and Parkinsonian syndromes, where imaging-derived effect maps are correlated with transcriptomic gradients to infer biological pathways and cell-type contributions underlying disease-specific patterns of neurodegeneration [5–7]. More recently, imaging transcriptomics has been applied to MSA in humans. In a landmark study, Chougar and colleagues [8] correlated regional atrophy maps derived from structural MRI in patients with MSA with gene expression data from the Allen Human Brain Atlas. Using multivariate statistical approaches, the authors identified molecular signatures enriched for oligodendrocyte-related and mitochondrial pathways that spatially aligned with MSA-specific atrophy patterns, providing new insights into the biological programs associated with selective regional vulnerability in this disorder. Importantly, this work demonstrated the feasibility and interpretive power of imaging transcriptomics in a rare and rapidly progressive neurodegenerative disease. At the same time, it is necessary to recognize the intrinsic limitations of imaging transcriptomics studies in clinical populations. The reliance on postmortem normative transcriptomic atlases precludes direct assessment of disease-state gene expression, and inter-individual variability, disease heterogeneity, and limited experimental control restrict mechanistic validation. In the context of MSA, these constraints make it difficult to disentangle whether imaging transcriptomic associations reflect primary drivers of neurodegeneration, downstream consequences, or compensatory responses. Moreover, validation of transcriptomic inferences at the tissue or cellular level is typically not feasible in human cohorts.

Experimental animal models offer a unique opportunity to address these limitations. The proteolipid protein (PLP)-αSyn transgenic mouse, which overexpresses human α-syn in oligodendrocytes [9], recapitulates key pathological features of MSA, including early oligodendroglial dysfunction and neurotrophic deficits, progressive α-syn aggregation, myelin dysfunction, gliosis, and selective neurodegeneration associated with progressive motor and non-motor deficits [10, 11]. Crucially, this model enables controlled, whole-brain MRI phenotyping, direct access to tissue for molecular validation, and integration of imaging findings with independently acquired transcriptomic data from the same disease context. Applying imaging transcriptomics in the PLP-αSyn mouse therefore serves several complementary purposes. First, it provides a preclinical testbed for evaluating whether imaging transcriptomic associations observed in human MSA are conserved across species and disease contexts. Second, it allows assessment of how different MRI modalities—such as volumetric and diffusion-based measures—map onto partially distinct molecular and cellular programs within a single, well-characterized model. Third, the availability of model-specific RNAseq data enables empirical evaluation of whether atlas-derived transcriptomic associations align with disease-relevant gene expression changes, thereby strengthening biological interpretability beyond what is possible in human studies alone. Importantly, the goal of imaging transcriptomics in the animal model is not to assert direct causality between gene expression and MRI phenotypes, but rather to identify molecular correlates of regional imaging vulnerability that can guide mechanistic hypotheses and targeted validation. Establishing these relationships at advanced stages of disease can provide a critical benchmark for interpreting earlier and subtler imaging changes, and will lay the groundwork for future studies focused on prodromal biomarkers and longitudinal progression.

In this study, we combine high-resolution ex-vivo structural and diffusion MRI with imaging-transcriptomics analysis in the PLP-αSyn mouse model of MSA. By integrating multimodal MRI phenotypes with more than 4,000 normative gene expression maps from the Allen Brain Atlas, followed by gene enrichment analyses to determine the associated biological processes and cell types [7, 12, 13] and validating key findings using model-specific RNAseq data, we aim to delineate the molecular and cellular programs associated with MRI-detected neurodegeneration and microstructural alterations in the MSA mouse model. This work extends imaging transcriptomics into a controlled experimental setting, complements recent human studies, and establishes a translatable framework for mechanistically informed interpretation of MRI biomarkers in synucleinopathies.

## RESULTS

### Multiparametric MRI identifies atrophy and microstructural alterations in the MSA mouse brains

Ex vivo deformation-based morphometry revealed widespread brain atrophy in PLP-αSyn mice compared with age-matched WT controls (**Figure 1a; Suppl. Fig. S2a**). Significant age × genotype interactions were detected in some regions; therefore, post hoc comparisons were performed. At 10 months, PLP-αSyn mice exhibited pronounced volume loss in the midbrain, hindbrain, and cerebellum, accompanied by ventricular enlargement, consistent with neurodegeneration. Conversely, localized volume increases were observed in regions of the cortex, amygdala, and striatum (**Suppl. Fig. S2a**). By 15 months, atrophy extended to additional brain areas, and no regions displayed volume gain (**Suppl. Fig. S2b**). This pattern was at least partly associated with an age-related volume increase in WT mice (**Suppl. Fig. S2c**).

**Figure 1.**
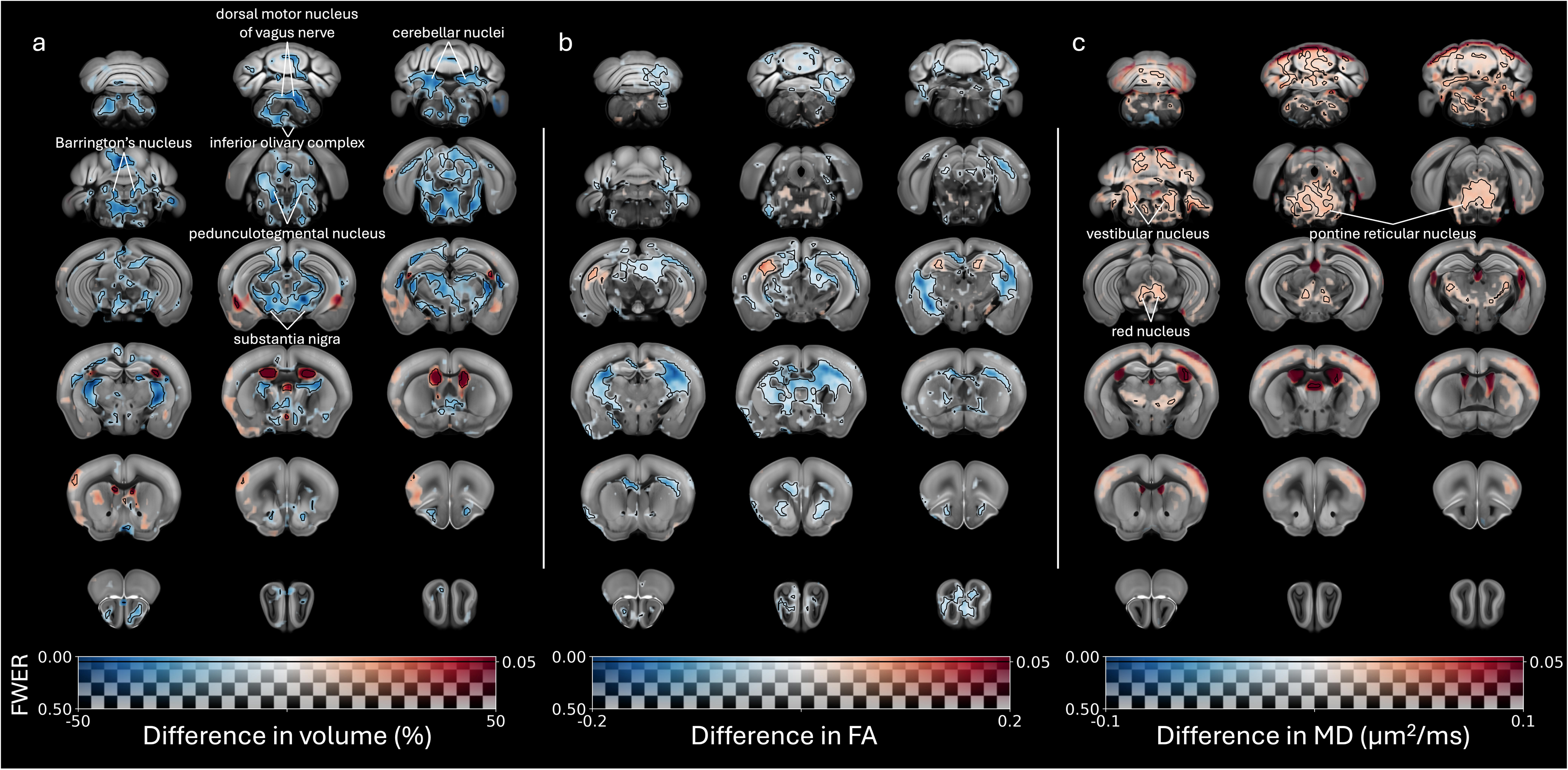
Multiparametric MRI phenotype of PLP-⍰Syn brains compared to wild-type mouse brains. Voxel-wise differences in a) volume, b) fractional anisotropy (FA), and c) mean diffusivity (MD) overlaid on the Allen Mouse Brain Common Coordinate Framework (CCFv3) template. The difference in group means (PLP-⍰Syn minus wild-type) is mapped to the color of the overlays, while the family-wise error rate (FWER) is mapped to the transparency of the overlays. The overlays are completely transparent where the FWER > 0.5, while areas where the FWER < 0.05 are contoured by black lines.

Given the limited group sizes and the absence of age or age × genotype effects in the diffusion metrics (FA and MD), subsequent analyses focused on the main genotype effect. Significant atrophy was identified in multiple motor-related structures, including the pons, extrapyramidal fiber tracts, medulla, pedunculopontine tegmental nucleus, and cerebellum (**Table 2**; **Figure 2a**). Non-motor regions implicated in sleep regulation, autonomic control, anxiety, and pain processing—such as the epithalamus, ventral tegmental area, raphe nuclei, insular cortex, reticular nucleus, and intralaminar thalamic nuclei—also showed reduced volume (**Suppl. Fig. S3a**).

**Figure 2.**
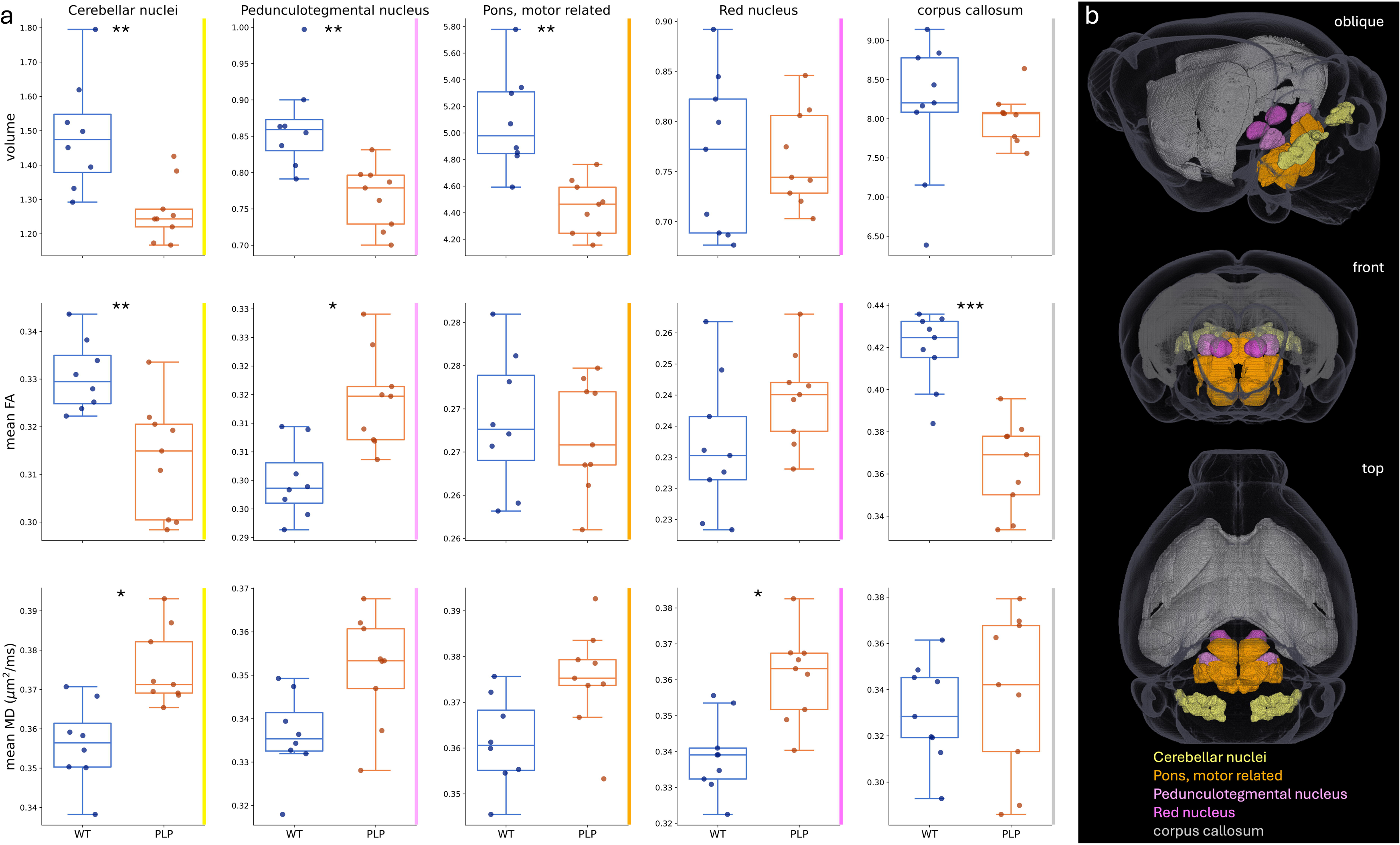
Most affected brain regions in PLP-⍰Syn mice according to atlas-based statistics of the MRI phenotype. a) Dot and box plots of regional volume, mean fractional anisotropy (FA), and mean mean diffusivity (MD) for select regions-of-interest (ROI). Two-tailed Mann-Whitney U tests, FDR-corrected p < 0.1*, 0.01**, 0.001***. b) 3-dimensional representation of the selected ROIs.

**Table 2.** Most affected brain regions according to atlas-based statistics comparing PLP-αSyn and wild-type mice.

| volume |  |  | FA |  |  | MD |  |  |
| --- | --- | --- | --- | --- | --- | --- | --- | --- |
| ROI | % difference in group medians | FDR-corrected p-value | ROI | % difference in group medians | FDR-corrected p-value | ROI | % difference in group medians | FDR-corrected p-value |
| epithalamus related | -8.89 | 0.004 | anterior commissure, olfactory limb | -12.32 | 5.10E-04 | Pons, behavioral state related | 4.86 | 0.05 |
| Perithalamus | -14.15 | 0.004 | optic nerve | -12.59 | 5.10E-04 | Cerebellar nuclei | 4.16 | 0.054 |
| Ventral tegmental area | -19.98 | 0.004 | thalamus related | -13.64 | 5.10E-04 | Red nucleus | 7.08 | 0.054 |
| Raphe nuclei | -14.7 | 0.004 | fornix system | -13.59 | 5.10E-04 | (Para)flocculus | 7.96 | 0.054 |
| Pons, sensory related | -11.73 | 0.004 | epithalamus related | -13.33 | 5.10E-04 | Cerebellar vermis | 8.84 | 0.064 |
| Pons, motor related | -10.33 | 0.004 | lateral ventricle | -19.32 | 5.10E-04 | Cerebellar hemispheres | 15.12 | 0.115 |
| medial lemniscus | -12.7 | 0.004 | third ventricle | -14.36 | 5.10E-04 | Inferior colliculus | 9.6 | 0.18 |
| extrapyramidal fiber systems | -11.97 | 0.004 | corpus callosum | -13.08 | 7.20E-04 | third ventricle | 19.3 | 0.18 |
| Insular cortex | 7.82 | 0.005 | fourth ventricle | -9.03 | 7.20E-04 | Pedunculotegmental nucleus | 5.36 | 0.18 |
| Reticular nucleus | -11.5 | 0.005 | choroid plexus | -18.24 | 7.20E-04 | arbor vitae | 3.87 | 0.18 |
| Medulla, motor related | -10.04 | 0.005 | stria terminalis | -13.1 | 0.001 | Pons, motor related | 4.07 | 0.217 |
| Intralaminar thalamic nuclei | -10.82 | 0.006 | Olfactory bulb | -6.93 | 0.003 | Ventral thalamus | 5.39 | 0.228 |
| Pedunculotegmental nucleus | -9.34 | 0.006 | Geniculate nucleus | -9.51 | 0.003 | facial nerve | 4.08 | 0.241 |
| cerebellar peduncles | -14.15 | 0.006 | Perithalamus | -9.83 | 0.003 | Raphe nuclei | 5.49 | 0.288 |
| internal capsule | -17.15 | 0.007 | Pretectal region | -13.58 | 0.003 | lateral ventricle | 36.01 | 0.292 |
| Pons, behavioral state related | -9.64 | 0.008 | Superior colliculus | -9.94 | 0.003 | oculomotor nerve | 5.18 | 0.319 |
| Cerebellar nuclei | -15.68 | 0.008 | arbor vitae | -9.3 | 0.003 | Medulla, sensory related | 2.72 | 0.319 |
| vestibulocochlear nerve | -13.28 | 0.008 | internal capsule | -9.75 | 0.004 | corticospinal tract | -2.03 | 0.319 |
| stria terminalis | -9.81 | 0.013 | Cerebellar hemispheres | -10.74 | 0.005 | Primary somatosensory cortex | 6.51 | 0.319 |
| Cuneiform nucleus | -6.82 | 0.016 | Cerebellar nuclei | -4.42 | 0.007 | Habenula | 8.99 | 0.319 |

FA was reduced across widespread white-matter tracts (**Table 2; Suppl. Fig. S3b**), including interhemispheric pathways (anterior commissure, corpus callosum), descending motor tracts (internal capsule, cerebellar peduncles), and fibers associated with non-motor systems (optic nerve, olfactory bulb, epithalamus, stria terminalis) (**Table 2**; **Figure 2b**). In contrast, MD was globally increased, with the most significant elevations in the brainstem and cerebellum (**Table 2**; **Figure 2c; Suppl. Fig. S3c**).

### Imaging transcriptomics highlights associations between MRI phenotypes and gene expression signatures of myelination, gliogenesis, neuroinflammation, and neural network regulation

To relate MRI changes to underlying molecular mechanisms, we performed imaging transcriptomic analyses, examining spatial correlations between MRI-derived t-statistic maps (volume, FA, MD) and normative gene expression patterns from the Allen Brain Atlas. We identified 3,214 genes significantly correlated with regional volume changes (p_adj_ < 0.05), 1,850 with FA, and 2,334 with MD (**Suppl. Tables S1–S3**). All correlations exceeded those expected by chance (p = 0 vs. 5,000 BrainSMASH surrogates). The top 30 volume- and top 55 FA-correlated genes were predominantly negatively associated with the corresponding MRI features, while among the top 60 MD-correlated genes, 20 showed positive associations (**Suppl. Tables S1–S3**).

Genes most strongly correlated with atrophy included *Arhgef10* (axon guidance, nerve conduction), *Cldn11* (axon ensheathment), *Tspan2* (oligodendrocyte differentiation), *Aspa* (N-acetylaspartate metabolism relevant to myelin synthesis), *Cnp* (myelinating cell RNA regulation), *Ugt8a* (galactocerebroside synthesis), and *Fa2h* (myelin sphingolipid biosynthesis). Top FA-correlated genes overlapped substantially, featuring *Cldn11*, *Tspan2*, *Cnp*, and *Fa2h*, alongside immune and proteostasis regulators (*Lpar1*, *CD9*, *App*, *Adamts4*, *Cryab*). MD-correlated genes included neuronal and metabolic markers such as *Pvalb* (inhibitory interneurons), *Scn1a* (neuronal excitability), *Uqcrh* (mitochondrial respiration), and *Abcd2* (energy metabolism).

To examine associations between PLP-driven human α-syn overexpression and MRI phenotypes, we correlated MRI t-statistic maps with regional *Plp1* expression (**Figure 3**). Regional volume loss and FA reductions showed negative correlations with *Plp1* expression, highlighting a spatial association between these MRI measures and oligodendrocyte-related transcriptional patterns. In contrast, MD showed no association with *Plp1* expression, consistent with its enrichment for neuronal and mitochondrial gene signatures and suggesting that diffusivity changes may reflect processes distinct from primary oligodendroglial pathology.

**Figure 3.**
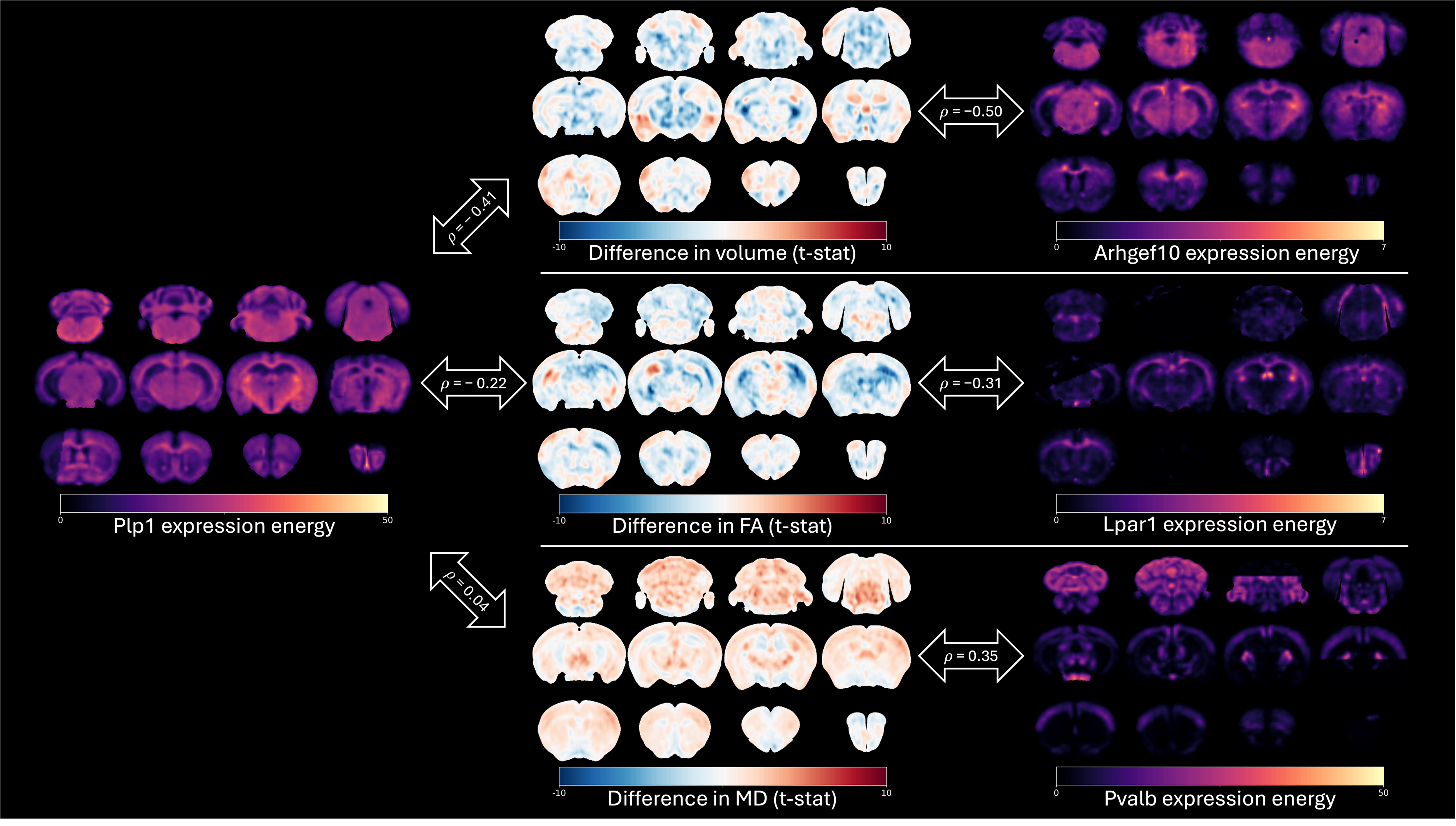
Spatial correlation between the PLP-⍰Syn MRI phenotype and Allen gene expression maps. Left: coronal slices of the Plp1 expression energy map from the Allen Mouse Brain Atlas. Middle: corresponding slices of t-statistic maps from voxel-wise comparisons of volume, fractional anisotropy (FA), and mean diffusivity (MD) in PLP-⍰Syn mice vs wild-types. Right: corresponding slices of gene expression energy maps that correlated most strongly with the t-statistic maps. The Spearman correlation coefficients (ρ) between the t-statistic and gene expression maps are shown in the double arrows.

Gene Ontology (GO) enrichment analysis of FA-correlated genes revealed significant (FDR < 0.05) overrepresentation of processes related to myelination, gliogenesis, and astrocyte and oligodendrocyte differentiation, all prominently featuring *Plp1* (**Figure 4; Suppl. Tables S4–S5**). By contrast, genes associated with MD changes were enriched for dendritic spine development, reflecting neuronal programs without implying mechanistic causality (**Suppl. Table S6**). The modality-specific associations of MRI maps with oligodendrocyte-related transcriptional programs argue against a purely circular effect and indicate that different MRI measures reflect multiple, complementary biological processes.

**Figure 4.**
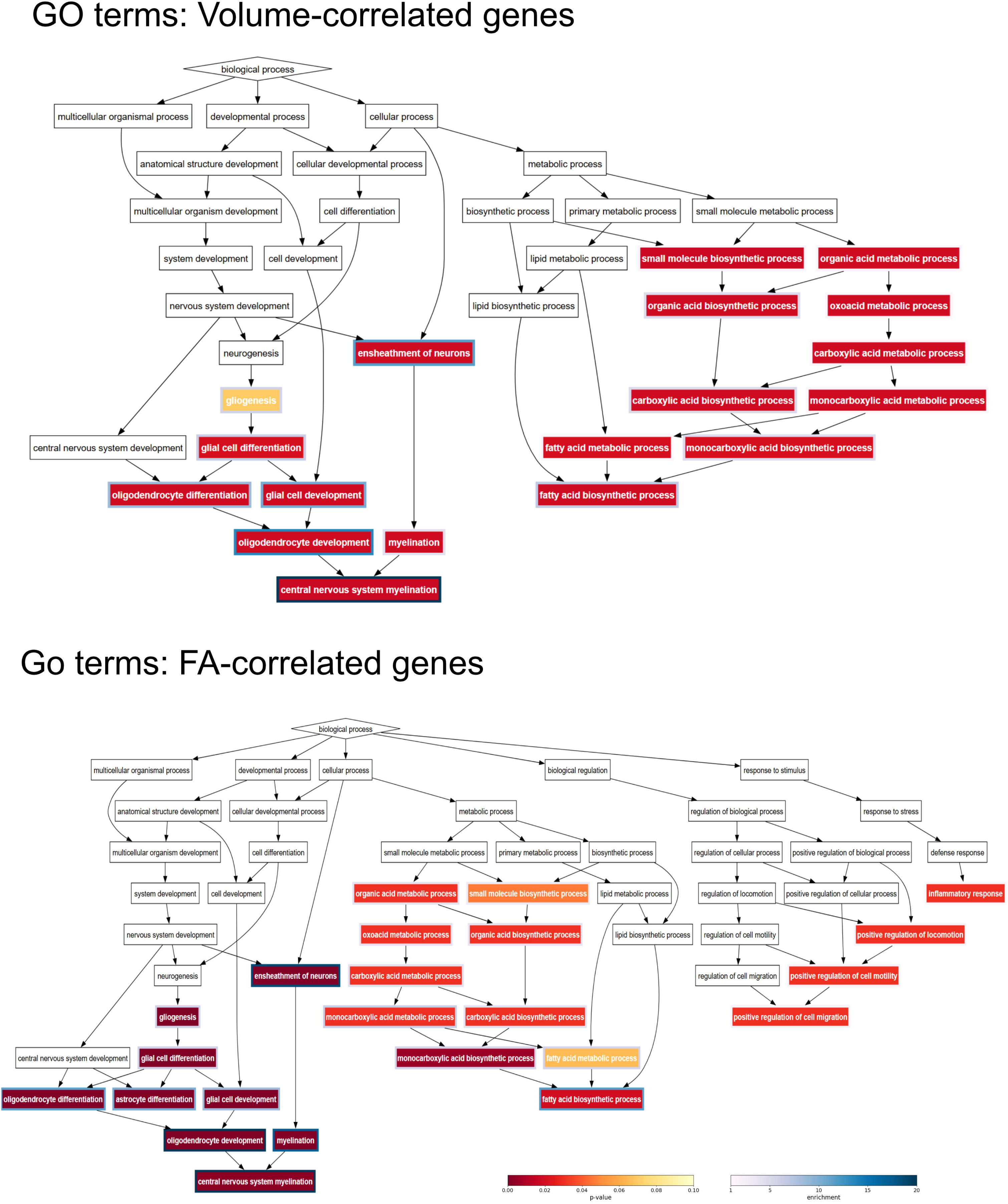
Enriched Gene Ontology terms among genes most spatially correlated with the PLP-⍰Syn MRI phenotype. Directed acyclic graphs of Gene Ontology (GO) biological process terms containing Plp1 that are significantly enriched (Benjamini-Hochberg-corrected p < 0.05) in the top of ranked lists of genes most correlated with PLP-⍰Syn changes in a) volume and b) fractional anisotropy (FA). For readability purposes, parent terms were included in the graph only if they are in the shortest path between a significantly enriched term and the root “biological process” term. The GO term nodes are colored if corrected p < 0.1, with the node outline color indicating the enrichment score. For a given GO term, enrichment = (b/n) / (B/N), where N is the total number of genes in the ranked list; B is the number of genes associated with the GO term; n is the size of the target set, i.e., the number of genes in the top of the ranked list that gives the minimum hypergeometric (mHG) score; and b is the number of genes in the target set that are associated with the GO term.

### Cell-type enrichment implicates oligodendrocytes, astrocytes, and specific neuronal subpopulations

Cell-type–specific enrichment analysis demonstrated that genes spatially correlated with regional volume loss were significantly enriched (FDR < 0.05) for oligodendrocyte markers in the cortex and cerebellum, and for cholinergic neurons of the spinal cord and brainstem (**Figure 5a; Suppl. Table S10**). Subthreshold enrichment (0.05 < p < 0.1) was observed for brainstem serotonergic neurons.

**Figure 5.**
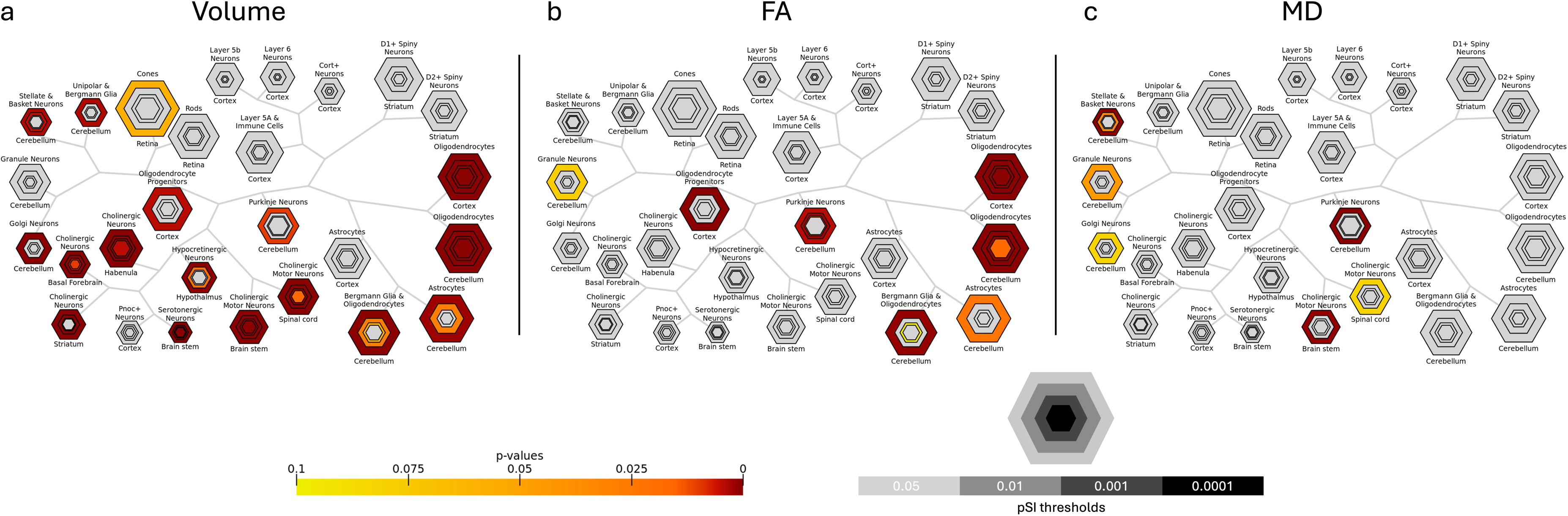
Enriched cell types in genes most spatially correlated with the PLP-⍰Syn MRI phenotype. Each cell type is represented by four concentric hexagons, the areas of which are proportional to the number of gene that are enriched in that cell type at four different Specificity Index (pSI) thresholds. The smallest, central hexagon represents the list of genes that are most specifically enriched in a particular cell type. The hexagons are grouped by brain region, indicated by the background color. Hexagons are colored if there is an enrichment (Benjamini-Hochberg-corrected p < 0.1) of genes associated with the corresponding cell-type in the ranked lists of genes most correlated with PLP-⍰Syn changes in a) volume, b) fractional anisotropy (FA), and c) mean diffusivity (MD).

Similarly, genes correlated with FA reduction were enriched for cortical and cerebellar oligodendrocytes, cerebellar stellate and basket cells, and Purkinje cells (pSI < 0.001), as well as cortical and cerebellar astrocytes and oligodendrocyte progenitors (pSI < 0.05) (**Figure 5b; Suppl. Table S11**). Subthreshold enrichment was observed for Bergmann glia and cerebellar granule cells. No cell type reached significance for genes associated with increased MD (**Figure 5c; Suppl. Table S12**), although mild enrichment was noted for cortical layer 6 pyramidal cells and prepronociceptin (PNOC)-expressing neurons.

### RNAseq supports the biological relevance of molecular signatures of neurodegeneration identified by imaging transcriptomics

To test if the genes we have identified indirectly to be associated with disorder-related imaging phenotypes are indeed dysregulated in the post-mortem brain tissue of the same model, we performed RNAseq and DESeq2 analysis on cerebellar tissue from PLP-αSyn mice [14], which we found significantly affected in MRI modalities (**Figure 1, 2**), and cross-referenced the 2613 differentially expressed genes (DEGs, FDR<0.05) with those correlated to MRI-derived maps. We identified 664 DEGs overlapping with volume-correlated genes, 441 with FA-correlated genes, and 510 with MD-correlated genes (**Suppl. Tables S7-S9**). The log2(FC) values of cerebellar DEGs were weakly but significantly correlated with FA-correlated genes (ρ =-0.106, p=0.026) and MD-correlated genes (ρ=-0.1895, p<0.0001) (**Figure 6**). Further enrichment analysis showed that the DEGs were significantly overrepresented (p=0.0376) among the genes whose normative spatial expression patterns correlated most strongly with volume changes in the PLP-αSyn mice (**Table 3**). The overlap between DEGs and MRI-correlated genes is consistent with the biological relevance of the imaging transcriptomics findings and highlights an association between MRI-detected alterations and molecular signatures of neurodegeneration in MSA.

**Figure 6.**
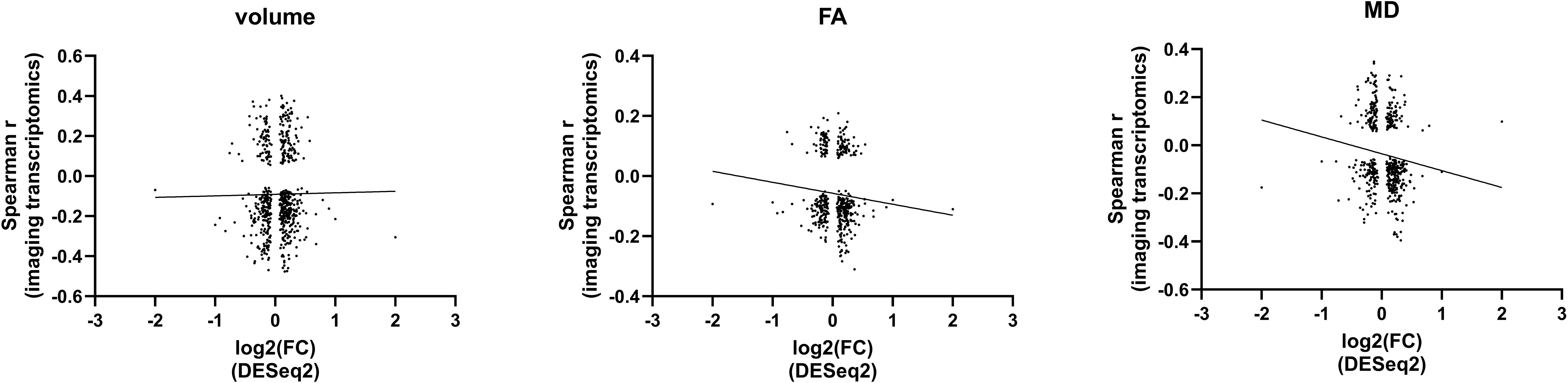
Spearman correlation analysis of DEGs in the cerebellum and MRI–linked genes identified by imaging transcriptomics. The Spearman correlation coefficients between genes’ normative spatial expression patterns (Allen Mouse Brain Atlas) and MRI-measured volume, fractional anisotropy (FA), and mean diffusivity (MD) changes in the PLP-αSyn brain are plotted against the genes’ log2(FC) from DESeq2 analysis of PLP-αSyn cerebellar tissue.

**Table 3.** Cerebellar DEG enrichment analysis.

| MRI parameter | N | B | b | n | enrichment | smHG | p-value |
| --- | --- | --- | --- | --- | --- | --- | --- |
| volume | 3494 | 758 | 146 | 504 | 1.33529128 | 1.96E-05 | 0.0376 |
| FA | 3494 | 758 | 9 | 18 | 2.30474934 | 0.00735212 | 0.3692 |
| MD | 3494 | 758 | 5 | 6 | 3.8412489 | 0.00234034 | 0.2398 |
| DEG: differentially expressed gene, N: total number of genes found in both the Allen Atlas and RNAseq datasets, B: number of DEGs in the N total genes, b: number of DEGs in the top n ranked genes, enrichment = $(b/n)/(B/N)$, smHG: minimum hypergeometric test statistic, FA: fractional anisotropy, MD: mean diffusivity | | | | | | | |

## DISCUSSION

Previous work has established the PLP-αSyn mouse as a valuable MSA model for investigating α-synuclein–driven mechanisms of neurodegeneration and testing disease-modifying therapies that mitigate α-syn aggregation, neuroinflammation, and neuronal loss [15–21]. Here, we provide the first comprehensive, whole-brain MRI characterization of this model compared with wild-type controls. High-resolution ex vivo structural MRI revealed a robust pattern of regional atrophy affecting white matter tracts, brainstem nuclei (medulla, pons, midbrain), cerebellum, thalamus, hypothalamus, and subiculum. The pattern of volume loss resembled that reported in MSA patients, with involvement of the hippocampus, pons/medulla, cerebellum, and pyramidal tracts (**Table 1**), supporting the translational relevance of MRI-detected neurodegeneration in the PLP-αSyn model. DTI revealed widespread microstructural abnormalities: FA was reduced across major white-matter tracts (e.g., corpus callosum, fornix, anterior commissure, striatothalamic fibres, and cerebellar peduncles), consistent with clinical MSA and suggestive of altered myelination and axon–glia interactions rather than overt demyelination. Focal FA increases in the hippocampus may reflect gliosis-related reorganisation or loss of crossing fibres. In contrast, MD was predominantly elevated in brainstem and cerebellum, consistent with tissue rarefaction and extracellular water increase. The modest overlap between FA and MD effects indicates that these metrics capture complementary aspects of pathology.

Applying an imaging transcriptomics framework to these multimodal MRI phenotypes, we observed that atrophy and microstructural alterations were systematically organized and aligned with transcriptional programs related to oligodendrocyte biology, energy metabolism, and neuroinflammatory processes. These findings broadly converge with human MSA imaging transcriptomics [8], despite differences in species, disease duration, imaging resolution, and experimental context, supporting the view that imaging–transcriptomic associations capture core molecular correlates of regional vulnerability.

A key extension beyond prior human studies is the integration of volumetric and diffusion-derived MRI measures. While structural atrophy reflects cumulative tissue loss, diffusion metrics such as FA and MD capture complementary microstructural features. FA and volume, but not MD, correlated with *Plp1* expression, linking these signatures to the oligodendroglial α-synucleinopathy, whereas MD aligned with neuronal and mitochondrial programs. Gene Ontology and cell-type enrichment analyses reinforce these modality-specific interpretations: oligodendrocytes and astrocytes were associated with FA and volume changes, whereas neuronal subtypes correlated with MD-related alterations. Importantly, different MRI phenotypes mapped onto partially distinct oligodendrocyte-associated transcriptional signatures, suggesting that regional vulnerability within the oligodendrocyte lineage—including myelination, metabolic support, and glial reactivity—may influence circuit-level susceptibility and pathological manifestation. In the controlled PLP-αSyn model, modality-specific signatures—previously unresolvable in clinical MSA—position diffusion MRI as a biologically informative complement to volumetric imaging for translational studies.

Imaging transcriptomics relies on normative gene expression atlases, which do not capture dynamic or region-specific transcriptional changes during neurodegeneration [5–7, 12]. In the PLP-αSyn model, we partially addressed this by comparing atlas-derived signatures with independent RNAseq data [14], revealing concordance that supports the biological relevance of the associations. While molecular validation was restricted to the cerebellum—a core site of pathology in both human MSA and the mouse model [1, 14]—the spatial alignment of MRI-detected alterations with known neurodegeneration sites reinforces the relevance of these imaging markers [10, 22–25].

We should point out that an immediate systematic histological confirmation was beyond the scope of this study but the model has been previously extensively histologically characterized [10, 22–27]. Among the limitations of this study is the reliance on ex vivo imaging in a modest cohort of female mice, limiting longitudinal assessment and raising the possibility of fixation-related bias. Future studies should incorporate longitudinal in vivo MRI with time-resolved transcriptomic profiling, and experimental manipulations such as conditional gene targeting or pharmacological interventions to establish causal links between cellular dysfunction and MRI phenotypes.

The primary value of applying imaging transcriptomics in the PLP-αSyn model lies in benchmarking and interpretability. By establishing how MRI phenotypes align with transcriptional programs in a controlled system, this work provides a reference framework for interpreting imaging findings in preclinical and clinical studies and enables direct comparison with human MSA data, strengthening confidence in shared disease mechanisms. Rather than replacing molecular or histological analyses, imaging transcriptomics adds value by integrating systems-level imaging with cellular biology, offering a scalable approach to contextualize MRI biomarkers and bridge the gap between macroscopic imaging readouts and cellular pathology in synucleinopathies.

In summary, our study is the first to apply whole-brain imaging to an MSA relevant mouse model and to show that its imaging phenotype parallels key features reported in human MSA. We also extend recent human work by applying imaging transcriptomics and we highlight both the strengths and limitations of this integrative approach. By combining multimodal MRI with atlas-based transcriptomic mapping and model-specific RNAseq, we provide a multiscale perspective on brain vulnerability in MSA and establish a foundation for future longitudinal and interventional studies to identify early imaging biomarkers and test mechanistic hypotheses.

## METHODS

### Animals

This study included nine female PLP-αSyn mice (four 10-month-old and five 15-month-old) and nine age-matched female wild-type (WT) mice (six 11-month-old and three 15-month-old). The animals were bred and aged in the Animal Facility of the Medical University Innsbruck under temperature-controlled pathogen-free conditions with a light/dark 12-hour cycle and free access to food and water. Mice were anesthetized with an overdose of thiopental and intracardially perfused with phosphate buffered saline (PBS pH_□7.4, Sigma-Aldrich, Vienna, Austria) followed by 4% paraformaldehyde (PFA, pH_□7.4, Sigma-Aldrich, Vienna, Austria). The heads were removed, postfixed in 4% paraformaldehyde for 24h at 4°C and stored in PBS with 0.05% sodium azide until scanning. All experiments were performed according to the ethical guidelines with the permission of the Austrian Federal Ministry of Science and Research (BMFWF-2022-0.899.398).

The sample size (n = 9 per group) was determined using the resource equation method, which ensures an efficient balance between statistical validity and ethical reduction principles [28]. In this design, the calculated error degrees of freedom (E = total number of animals – total number of groups = 18 − 2 = 16) fall within the acceptable range of 10 ≤ E ≤ 20, indicating adequate power for detecting biologically meaningful effects while minimizing unnecessary animal use. The chosen group size is consistent with previous preclinical MRI studies in transgenic models of neurodegeneration. Only female mice were included to reduce variability associated with sex differences in brain structure.

### Magnetic resonance imaging

#### Image acquisition

Skulls were placed four at a time in a 50-ml Falcon tube, separated and secured by a custom 3D-printed holder, and immersed in perfluoropolyether (Galden®, Solvay). These were scanned in a 9.4T Bruker BioSpec 94/20 preclinical MRI scanner with a 39-mm radio-frequency coil to acquire T2-weighted (T2w) structural images and diffusion tensor imaging (DTI) data as previously described [29]. In brief, T2w images with 100-micron isotropic voxel size were acquired with a 3D fast spin echo sequence, and DTI images with 200-micron isotropic voxel size were acquired with a 2D multi-slice pulsed gradient spin echo sequence (4 b0 images, 30 diffusion directions, b=1500 s/mm2).

#### Image processing and analysis

Diffusion tensor fitting was performed using the dtifit command in FMRIB Software Library (FSL version 5.0.10). Multi-modal study-specific templates were created from the resulting fractional anisotropy (FA) and mean diffusivity (MD) maps and the T2w images using the antsMultivariateTemplateConstruction2.sh script included in Advanced Normalization Tools (ANTs version 2.2.0). Template and individual subject brain masks were created by multiplying the reciprocal of the MD map and the T2w image (fslmaths) and passing this image product to the Rapid Automatic Tissue Segmentation (RATS) tool for skull-stripping of rodent brain MRI [30, 31]. Masked subject images were registered to the masked templates using antsRegistration to perform sequential rigid-body, affine, and diffeomorphic SyN multi-modal registrations with equal weight given to the FA, MD, and T2w images. To quantify inter-subject voxel-level variations in local brain volume, maps of the Jacobian determinants of the composite transformations were computed using the ANTs CreateJacobianDeterminantImage command. The registered FA and MD maps were smoothed with a full-width half-maximum of 0.2 mm using the 3dBlurInMask command in the Analysis of Functional NeuroImages (AFNI) software suite (version 17.2.17). Univariate, 2×2 ANOVAs were performed on the log-transformed Jacobian determinant maps and the smoothed DTI maps to test for the effects of age (old vs. middle age) and genotype (PLP-αSyn vs. WT) on volume, FA, and MD. This was done using FSL’s randomise tool for nonparametric permutation inference to perform 10,000 permutations, threshold-free cluster enhancement, and family-wise error correction.

For region-of-interest (ROI)-based analysis, the masked T2w study template was registered to the Allen Mouse Brain Common Coordinate Framework (CCFv3) using antsRegistration as described above. The CCFv3 atlas labels were merged into 87 ROIs and warped to study template space. ROI volumes were computed by taking the sum of all Jacobian determinant values within each ROI and multiplying by the volume of one voxel. Mann-Whitney U (MWU) tests were performed to compare ROI volumes and mean FA and MD values from the PLP-αSyn and WT groups. The Benjamini-Hochberg procedure was used to correct for multiple comparisons (87 ROIs) for each MRI parameter.

#### Spatial gene enrichment analysis

Given the limited sample size, all mice (10-15 months of age) were pooled and only the main effects of genotype were considered for spatial gene enrichment analysis. Only the coronal dataset of the Allen *in situ* hybridization data was used because, although it contains far fewer genes than the sagittal dataset, it has superior brain coverage and quality. After excluding gene expression energy maps that had missing data in greater than 20% of voxels within the brain, the final number of maps included in the analysis was 4071 (some genes have duplicates, which were treated separately and all kept for correlation analysis).

#### Imaging-transcriptomic correlation

The volume, FA, and MD t-statistic maps generated by randomise were warped to CCFv3 space and down-sampled to 200-micron isotropic voxel size to match the gene expression energy maps. The Spearman correlation coefficient (ρ) was calculated between each MRI t-statistic map and each gene expression energy map.

To calculate non-parametric p-values for the ρ values, the BrainSMASH Python package [32] was used to generate 5000 surrogate maps that match the spatial autocorrelation of each of the volume, FA, and MD t-statistic maps. The Spearman correlation coefficient was computed between each surrogate map and each gene expression energy map to build null distributions from which p-values were computed (**Suppl. Fig. S1**).

#### Gene set enrichment analysis

Two separate gene set enrichment analyses were performed using two categories of gene sets: gene sets associated with the Gene Ontology (GO) biological process terms, and cell-type specific expression (CSE) gene sets curated by Dougherty et al. [33].

For each of 27 different cell types, the authors identified genes that are enriched in that cell population. Further, they truncated these gene lists at four Specificity Index thresholds (pSI) (a smaller pSI corresponds to a more exclusive list of genes that are more specific to a particular cell type) (http://doughertytools.wustl.edu/CSEAtool.html). Two cell types—cone and rod cells—were deemed irrelevant to this study and excluded.

For each type of analysis, only genes found in both the filtered Allen coronal dataset and the GO or CSE gene sets were included, resulting in 3606 and 3646 expression maps of 3385 and 3422 unique genes, respectively. The GO biological process terms were filtered by removing those with fewer than 10 or more than 200 remaining genes, à la Ritchie [34]. Furthermore, terms with identical gene sets (exactly equal) to another term were discarded, leaving a final count of 2393 GO biological process terms. The curation of the GO and CSE gene sets were performed with custom Python scripts.

For each MRI parameter, genes were ranked by FDR-corrected p-value and then by the absolute value of ρ. For genes with duplicates, only the highest ranked duplicate was kept for gene set enrichment analysis. The minimum hypergeometric (mHG) score [35] was used to quantify the enrichment of GO and CSE gene sets at the top of these ranked lists. To account for propagating effects of spatial autocorrelation into the enrichment analysis, a null distribution of mHG scores for each MRI parameter and gene set pair was calculated from the BrainSMASH-derived surrogate maps. For each observed mHG score *s*, a p-value was calculated as the fraction of mHG scores in the corresponding null distribution *s’* < *s*. To control the false discovery rate in each MRI parameter, the Benjamini-Hochberg procedure was performed across: 1) all 2393 GO gene sets, 2) the 42 GO gene sets that contain Plp1, and 3) the 25 CSE gene sets for each pSI level.

#### Imaging transcriptomics and RNAseq correlation

To consolidate the relevance of the disease-associated gene maps defined by imaging transcriptomics, we identified the significantly correlated genes to volume, FA and MD (p<0.05), which are found significantly dysregulated in the PLP-αSyn cerebellum by RNAseq and DESeq2 analysis (p_adj_>0.05,[14]). We calculated the Spearman correlation of the Spearman ρ values from the imaging transcriptomics and the Log2(FC) values from the DESeq2 analysis applying GraphPad Prism 10.5.0 software. In addition, we performed the enrichment analysis described above using the significantly dysregulated genes as a gene set. Of the 3494 genes found in both the Allen Atlas and RNAseq datasets, 758 genes were differentially expressed and considered in this enrichment analysis.

## Supplementary material

**Additional file 1**

Title: Supplementary Figures: Fig S1 to S3.

Format: .pdf

**Additional file 2**

Title: Supplementary Tables: Tables S1 to S12.

Format: .xls

Captions:

**Suppl Table S1**. Genes significantly correlated with regional volume changes.

**Suppl Table S2**. Genes significantly correlated with regional FA changes.

**Suppl Table S3**. Genes significantly correlated with regional MD changes.

**Suppl Table S4**. GO enrichment analysis of genes correlated with volume changes.

**Suppl Table S5**. GO enrichment analysis of genes correlated with FA changes.

**Suppl Table S6**. GO enrichment analysis of genes correlated with MD changes.

**Suppl Table S7**. DEGs in RNAseq of cerebellum (p<0.05, Log2(FC)) and significantly correlated with volume changes (p<0.05, Spearman r).

**Suppl Table S8**. DEGs in RNAseq of cerebellum (p<0.05, Log2(FC)) and significantly correlated with FA changes (p<0.05, Spearman r).

**Suppl Table S9**. DEGs in RNAseq of cerebellum (p<0.05, Log2(FC)) and significantly correlated with MD changes (p<0.05, Spearman r).

**Suppl Table S10**. Cell-type–specific enrichment analysis of genes correlated with volume changes

**Suppl Table S11**. Cell-type–specific enrichment analysis of genes correlated with FA changes

**Suppl Table S12**. Cell-type–specific enrichment analysis of genes correlated with MD changes

## Availability of data and materials

All data generated or analysed during this study are included in this published article and its supplementary information files.

## Competing interests

EK, declares no competing interests

DC, declares no competing interests

DM, declares no competing interests

CS, declares no competing interests

EK, declares no competing interests

AHG, received grant support from the Austrian Science Fund (FWF) and the Tyrolian Science Fund (TWF) outside of the submitted work.

FK, received personal fees from AbbVie, Bial, Ionis Pharmaceuticals, Koneksa, Merz, Österreichische Apotheker-Verlagsgesellschaft, Sanofi, Takeda Pharmaceuticals, and Teva and his institution has ongoing grant support from the Austrian Science Fund (FWF), the National Institutes of Health and the Michael J. Fox Foundation, outside of the submitted work.

LM, declares no competing interests

NS, received grant support from the Medical University of Innsbruck and The Michael J. Fox Foundation, outside of the submitted work; Director-at-large, Board of Directors and Chair of the Research Steering Council of Mission MSA, USA; Advisory board, Karl Golser Foundation, Italy; Advisory services for IONIS, Mitsubishi Pharma

## Funding

DM is supported by NIHR Maudsley Biomedical Research Centre, South London and Maudsley NHS Trust.

## Authors’ contributions

EK has contributed to the methodology, data analysis of the study and drafting parts of the manuscript

DC has contributed to the conceptualization of the study, data interpretation, and drafting parts of the manuscript

DM has contributed to the methodology of the study CS has contributed to the methodology of the study

EK has contributed by summarizing clinical MRI/DTI studies in MSA

AHG has contributed to data analysis and tissue preparation

FK has contributed to the interpretation of the clinical relevance of the data

LM has contributed to the conceptualization of the study

NS has contributed to the conceptualization of the study, analysis of the data, data interpretation, and drafting parts of the manuscript

All authors have reviewed and approved the final version of the manuscript.

## Notes

### Summary of Updates

The revision includes updates to the Introduction and Discussion to more clearly present and contextualize the significance of imaging transcriptomics methodology

