## Supplementary material for "Multiparametric MRI and imaging transcriptomics reveal molecular and cellular correlates of neurodegeneration in experimental parkinsonism": Fig. S1-S3

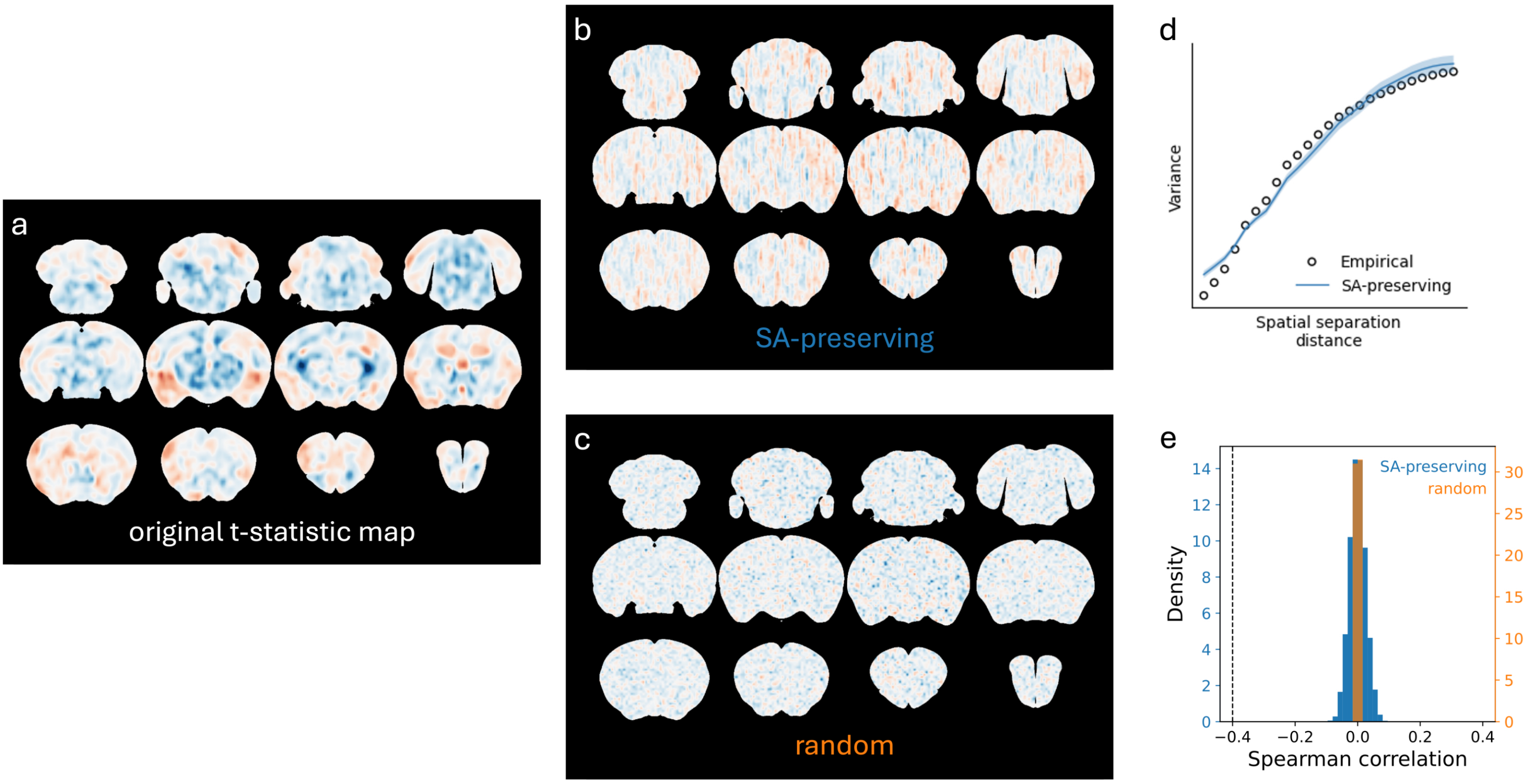


**Suppl. Fig.S1.** Spatial-autocorrelation-preserving surrogate brain maps created by the BrainSMASH toolbox. a) Coronal slices of the empirical t-statistic map resulting from the voxel-wise comparison of log-Jacobians between PLP-αSyn and WT mice. b) An example spatial autocorrelation (SA)-preserving surrogate of the t-statistic map. c) An example surrogate map created by simple random reshuffling of voxels. d) A variogram demonstrating similar spatial autocorrelation structure between the empirical t-statistic map and the SA-preserving surrogate maps. e) ull distributions of Spearman correlation coefficients between the Plp1 gene expression energy map and 5000 SA-preserving or randomly permuted surrogate t-statistic maps.

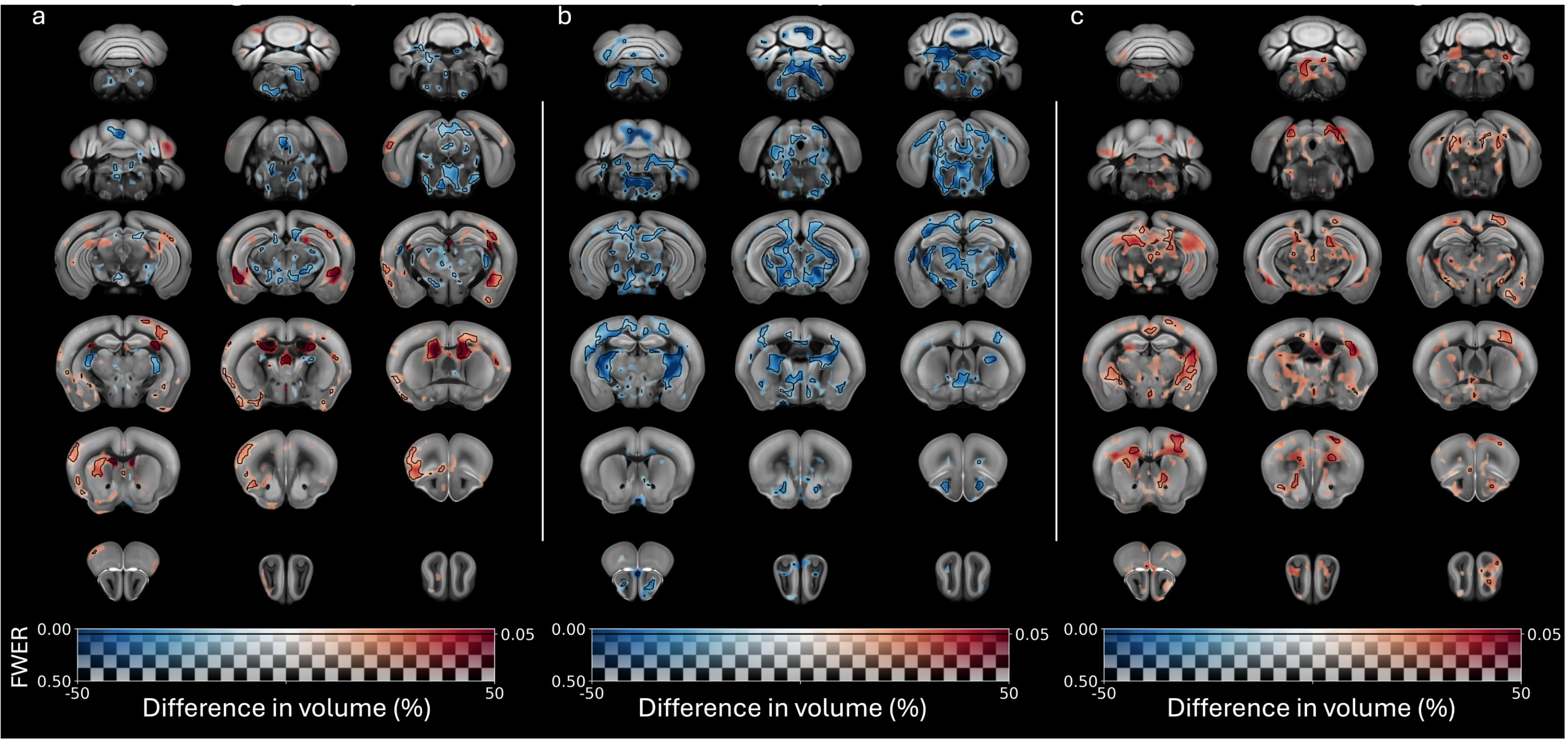


**Suppl. Fig. S2.** Post hoc voxel-wise comparisons of local brain volumes. Voxel-wise differences in volume between a) 10/11-month old PLP-αSyn / wild-type (WT) mice, b) 15-month old PLP-αSyn / WT mice, and c) WT 15- versus 11-month old mice. The difference in group means is mapped to the color of the overlays, while the family-wise error rate (FWER) is mapped to the transparency of the overlays. The overlays are completely transparent where the FWER > 0.5, while areas where the FWER < 0.05 are contoured by black lines.

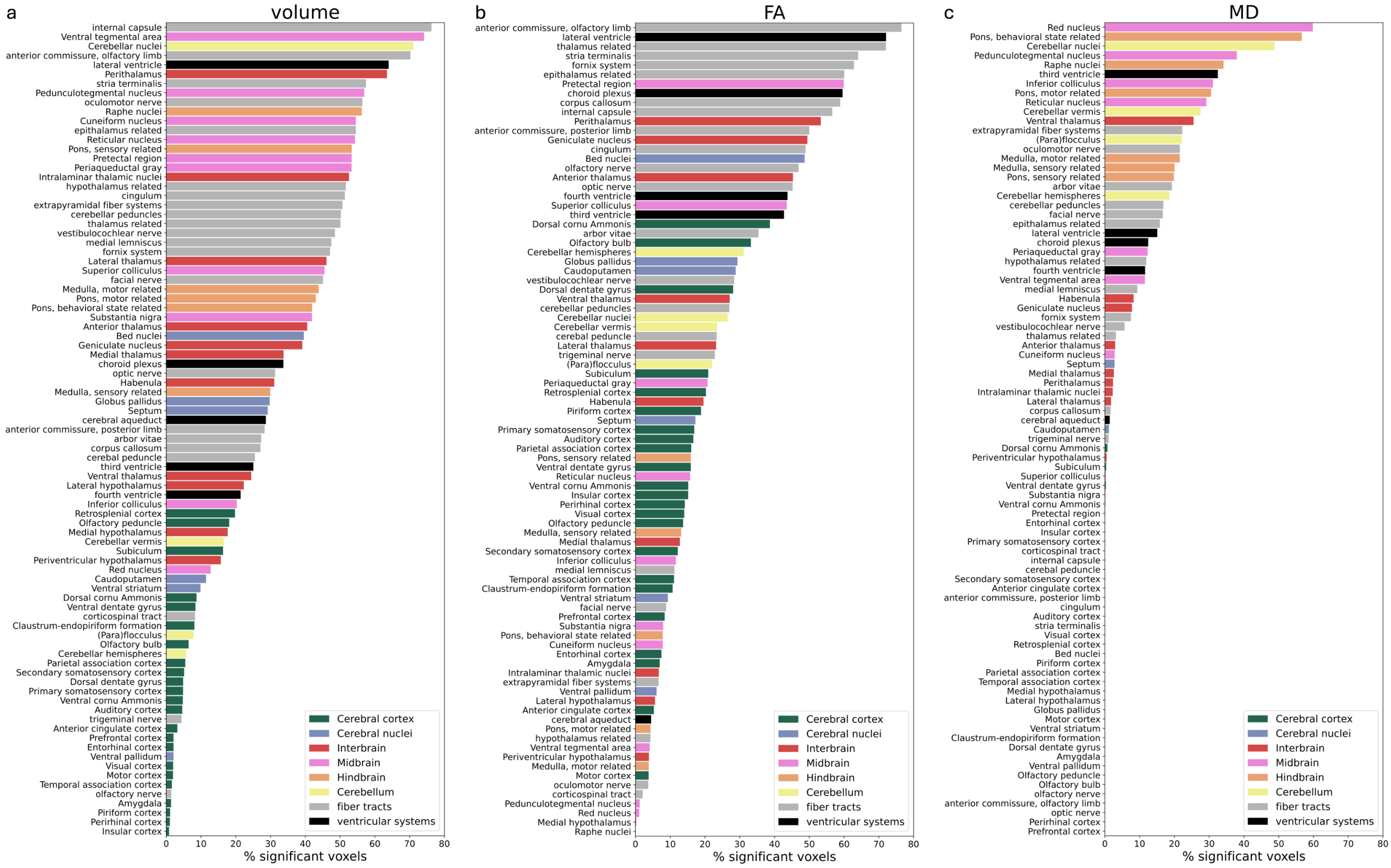


**Suppl. Fig. S3.** Region-by-region breakdown of voxel-wise differences between PLP-αSyn and wild-type mice. Bar graphs showing the percentage of voxels within each atlas region that had a significant (family-wise error rate < 0.05) difference in a) volume, b) fractional anisotropy (FA), and c) mean diffusivity (MD). Regions are color-coded based on the Allen Mouse Brain Atlas hierarchy.
